# Sex-dependent influence of major histocompatibility complex diversity on fitness in a social mammal

**DOI:** 10.1101/2024.12.18.629201

**Authors:** Nadine Schubert, Hazel J. Nichols, Francis Mwanguhya, Robert Businge, Solomon Kyambulima, Kenneth Mwesige, Joseph I. Hoffman, Michael A. Cant, Jamie C. Winternitz

**Affiliations:** Department of Animal Behavior, Bielefeld University, Bielefeld, Germany; Department of Biosciences, Swansea University, Singleton Campus, Abertawe, UK, SA2 8PP, and also Department of Animal Behavior, Bielefeld University, Bielefeld, Germany; Banded MongooseResearch Project, Queen Elizabeth National Park, Kasese District, Rubirizi, Uganda; Department of Evolutionary Population Genetics, Faculty of Biology, Bielefeld University, 33501 Bielefeld, Germany, and Center for Biotechnology (CeBiTec), Faculty of Biology, Bielefeld University, 33615 Bielefeld, Germany, and British Antarctic Survey, High Cross, Madingley Road, Cambridge CB3 OET, UK, and Joint Institute for Individualisation in a Changing Environment (JICE), Bielefeld University and University of Münster, Bielefeld, Germany; Centre for Ecology and Conservation, University of Exeter, Penryn Campus, Penryn TR10 9FE, UK; Department of Evolutionary Immunogenomics, Institute for Animal Cell and Systems Biology, University of Hamburg, Martin-Luther-King-Platz 3, 20146 Hamburg, Germany, and also Department of Animal Behavior, Bielefeld University, Bielefeld, Germany

## Abstract

Parasite infections affect males and females differently across a wide range of species, often due to differences in immune responses. Generally, females tend to have stronger immune defenses and lower parasite loads than males. The major histocompatibility complex (MHC) plays a crucial role in the adaptive immune response, and extensive research has explored how variation in this region influences infection and fitness outcomes. However, studies of sex-specific relationships between MHC variation and infection are scarce, perhaps because MHC genes are located on the autosomes, which are shared by both sexes. Here, we provide evidence of sexually antagonistic selection in a wild, group-living mammal—the banded mongoose. Using genetic and life history data collected from over 300 individuals across 25 years, we found that particularly MHC class I (MHC-I) but also MHC class II (MHC-II) diversity influence lifetime reproductive success differently in males and females. Specifically, higher MHC diversity is linked to increased fitness in males but decreased fitness in females. Furthermore, MHC diversity did not differ between the sexes, indicating an unresolved genetic sexual conflict. Our findings demonstrate that sexually antagonistic selection acts on the MHC and may operate across both MHC classes but differently. This study contributes to the growing body of evidence that sex is a significant factor in shaping host immunity and fitness.

## Introduction

Fitness proxies including longevity and reproductive success can be strongly impacted by parasites (e.g. Leivesley et al. 2019). Since parasites, broadly defined to include macroparasites (e.g. helminths, arthropods and protozoa) and microbial pathogens (i.e. viruses, bacteria, and fungi), are detected and fought by the immune system, variation in the effectiveness of the immune response affects parasite load (reviewed in Klein and Flanagan 2016) and can thereby influence individual fitness by determining an individual’s infection status (e.g. Marzal et al. 2005). This variation in the immune response results in some individuals being more susceptible to infection than others. For example, sex, genetic heterozygosity and certain genes have been identified as factors influencing parasite loads (e.g. Alexander and Stimson 1988; Zuk and McKean 1996; Meyer-Lucht and Sommer 2005; Kloch et al. 2010; Mitchell et al. 2017; Budischak et al. 2023).

A crucial component of the adaptive immune response against parasites is the major histocompatibility complex (MHC). The MHC is a gene complex that is present in virtually all jawed vertebrates (Kaufman 2018) and comprises one of the most polymorphic regions of the vertebrate genome (Klein 1986). It encodes cell-surface glycoproteins that are crucial for the initiation of the adaptive immune response (Bjorkman et al. 1987). These MHC molecules bind and present self and foreign peptides to professional immune cells which can cause further activation of the immune response (Knapp 2005). The MHC is subdivided into two classes, MHC class I (MHC-I) and MHC class II (MHC-II), where MHC-I molecules are present on nearly all nucleated cells and mainly present peptides originating from intracellular sources (e.g. self-derived peptides and peptides originating from viruses or other pathogens that have entered the cell), while MHC-II molecules are present on professional antigen-presenting cells (APCs, including macrophages, B cells and dendritic cells), and present peptides originating mostly from exogenous sources (e.g. derived from bacteria or macroparasite particles that have been engulfed by the cell) (Neefjes et al. 2011). Thus, the MHC has the potential to affect the efficacy of immune responses aimed at parasites and thereby influence individual fitness.

As each MHC molecule variant can only bind a subset of all available peptides that can stimulate an immune response (Rammensee et al. 1999), the diversity or composition of the MHC determines the repertoire of pathogens that can be eliminated. The maximal diversity hypothesis proposes that individuals with higher allelic diversity at the MHC should respond more effectively to pathogens through increased recognition and elimination (Doherty and Zinkernagel 1975), leading to lower parasite loads. This negative relationship between MHC allelic diversity and infection intensity (Kloch et al. 2010; Radwan et al. 2012; Biedrzycka et al. 2018) coupled with a positive relationship between MHC diversity and fitness has been observed in several vertebrate species (Bonneaud et al. 2004; Thoß et al. 2011; Agudo et al. 2012; Schaschl et al. 2012; Pineaux et al. 2020). An alternative, called the optimal diversity hypothesis (Milinski 2006), proposes that an intermediate number of MHC alleles enables optimal protection against parasites, due to a tradeoff between increased peptide binding and T-cell depletion with higher numbers of individual MHC allelic variants (Nowak et al. 1992). However, there is little direct immunological evidence for this hypothesis (Migalska et al. 2019; Krishna et al. 2020) and a number of indirect studies investigating the link between MHC diversity and parasite load on fitness have failed to support either the maximal or the optimal diversity hypothesis (Harf and Sommer 2005; Meyer-Lucht and Sommer 2005; Radwan et al. 2012; Westerdahl et al. 2012; Sepil, Lachish, Hinks, et al. 2013; Sepil, Lachish, and Sheldon 2013). Hence, the mechanisms linking MHC diversity, parasite load and fitness remain elusive.

Immune responses against parasites have been observed to differ between the sexes, with females having lower parasite loads compared to males (Zuk and McKean 1996). This difference has been mainly attributed to differences in sex hormones (reviewed in Roved et al. 2017). Briefly, testosterone in males suppresses, while estrogen and progesterone in females enhance immune responses, resulting in higher parasite loads in males and increased susceptibility to autoimmune diseases in females. Thus, sexually antagonistic selection is predicted to result in different optimal allele numbers for the sexes, with a higher optimum predicted for males compared to females (Roved et al. 2017). In line with this, Roved et al. (2018) investigated the MHC-I in great reed warblers and, despite expecting MHC-II to be a the main target of sexually antagonistic selection due to its stronger antagonistic regulation through sex hormones (Roved et al. 2017), found that males with higher MHC-I diversity had greater reproductive success, whereas females with lower MHC-I diversity had greater reproductive success. There was no difference in MHC-I diversity between the sexes, indicating that neither of the sexes reached its MHC-I diversity optimum, suggesting an unresolved sexual conflict. Nevertheless, studies investigating sexual conflict in MHC optima are rare, often focus on a single MHC class, and they do not cover all of the residues responsible for peptide binding (PBR) and hence fail to capture all of the relevant functional diversity.

A long-term (25+ year) study of banded mongooses (*Mungos mungo*) provides an ideal system for studying the impact of MHC variation of fitness in a wild vertebrate population. Banded mongooses are small (∼1.5 kg) mammals that live in groups consisting of 10-40 adult individuals (Cant et al. 2016). In contrast to most cooperative breeders, reproductive skew is relatively low, with adults of both sexes regularly reproducing. Breeding is synchronized within each social group, with multiple females giving birth together, usually on the same night (Hodge et al. 2011). The resulting pups are raised communally by the group; Unusually, there is no evidence that mothers can recognize their own pups or vice versa (Marshall et al. 2021). In our study population, individuals show limited dispersal and frequently breed within their natal group, resulting in close relatives often being among the pool of potential mates (Nichols et al. 2014). This, combined with the difficulty of recognizing relatives within groups, results in high variance in inbreeding. Wells et al. (2018) used a 9-generation deep pedigree to reveal that mild inbreeding is widespread, with many individuals (66.4%) having non-zero inbreeding coefficients (mean pedigree inbreeding coefficient = 0.058), and an unusually high proportion of individuals (7.1%) being closely inbred (inbreeding coefficient > 0.25), resulting from full sibling or parent-offspring matings. This inbreeding has fitness consequences for reproductive success in males (Wells et al. 2018) and homozygosity increases susceptibility to parasite infections (Mitchell et al. 2017). Nevertheless, the negative effects of inbreeding are to some extent buffered by social care (Wells et al. 2020), while evidence has also been found for inbreeding avoidance, mediated by extra-group mating (Nichols et al. 2015) and non-random breeding within social groups (Sanderson et al. 2015; Khera et al. 2021). This targeted mate choice may facilitate the maintenance of relatively high MHC diversity despite frequent inbreeding (Schubert et al. 2024). The availability of detailed longitudinal life-history data for our study population, together with high variance in inbreeding, make the banded mongoose an ideal study system for investigating an effect of MHC variation on fitness components in a wild mammal population.

Here, we genotyped a total of 465 individuals for MHC-I exons 2 and 3 and MHC-II DRB exon 2 to capture the antigen-binding sites of both MHC molecules. We first investigated associations between neutral genetic diversity and MHC diversity, hypothesizing that they would be at least weakly positively correlated. We then used the MHC sequences to estimate three different measures of MHC functional diversity, which were combined with long-term life history data to investigate the influence of MHC variation on three fitness measures: (i) pup survival, (ii) adult survival, and (iii) lifetime reproductive success (LRS). We hypothesized that individuals with higher MHC diversity should have increased fitness, but that the effects of the MHC could be non-linear and sex-specific.

## Materials and Methods

### 2.1 Study site and field data collection

Data used in this study were collected on a wild population of banded mongooses inhabiting Queen Elizabeth National Park in Uganda (0°12’S, 27°54E’), which has been subjected to regular and systematic behavioral, life-history and genetic data collection for over 25 years. The study area consists of approximately 10 km² savannah including the Mweya peninsula and the surrounding area and the population is made up of 10-12 social groups at any one time, corresponding to approximately 250 individuals. All individuals are habituated to human observation at <10m (usually <5m), allowing us to collect detailed behavioral and life history data from each social group, which are visited every 1-4 days. To enable reliable location of our study animals, one or two adult individuals per group were fitted with a 27g radio collar (<2% of an individual’s body mass, Sirtrack Ltd., New Zealand) with a 20cm whip antenna (Biotrack Ltd., UK). Individual mongooses can be visually identified in the field through (1) dye patterns in the fur created using commercial hair dye (L’Oreal, UK) for individuals less than 6 months of age, and (2) shaved fur patterns or (3) color-coded collars made from plastic for adults that will not grow any further. Collars and shave patterns were maintained during trapping events regularly taking place every 3-6 months as described by Cant (2000), Hodge (2007), and Jordan et al. (2010). Individuals were given either an individual tattoo or a subcutaneous pit tag under anesthetic (TAG-P-122IJ, Wyre Micro Design Ltd., UK) during their first capture to allow permanent identification. Moreover, a 2mm tail tip tissue sample was taken for genetic analysis. Individual age was calculated based on the time between the mothers’ parturition date and, since deaths were not usually directly observed, the date of the disappearance of each individual from their group. Death could be distinguished from dispersal, since banded mongooses do not disperse individually, but in small groups (Cant et al. 2002) following a period of aggression from the rest of the group (Thompson et al. 2016). Lifespan was calculated as the timespan in days between the birth and death date. Data on daily rainfall in mm was obtained from the Mweya meteorological station, which is situated in the center of the study site. Gaps in the meteorological data, which made up less than 5% of the relevant timespan, were imputed using Kalman Smoothing (Hyndman and Khandakar 2008).

### 2.2 Ethical statement

Research was conducted under approval of the Uganda National Council for Science and Technology, the Uganda Wildlife Authority and the Ethical Review Committee of the University of Exeter. All research procedures adhered to the ASAB Guidelines for the Treatment of Animals in Behavioural Research and Teaching (ASAB Ethical Committee and ABS Animal Care Committee 2022).

### 2.3 Genetic analyses

#### 2.3.1 DNA extraction and microsatellite genotyping

We extracted DNA from 465 individuals using the Qiagen® DNeasy blood and tissue kit according to the manufacturers protocol. These individuals were genotyped at 35-43 microsatellite loci as described in detail by Sanderson et al. (2015).

#### 2.3.2 Estimation of heterozygosity

Microsatellite genotypes were used to estimate individual standardized multilocus heterozygosity (sMLH) using the R package InbreedR (Stoffel et al. 2016). Outliers with sMLH values of zero or samples for which microsatellite amplification was successful for too few loci were removed.

#### 2.3.3 MHC genotyping

MHC genotyping and data processing was carried out according to the protocol described in Schubert et al. (2024). In short, we amplified the MHC-I exon 2 using primers (F: CCACTCCCTGAGGTATTTCTACACC, R: CTCACCGGCCTCGCTCTG) based on the sequences published by Yukhi & O’Brien (1990), and the MHC-I exon 3 (F: GGTCACACAGCATCCAGAGA, R: GCTGCAGCGTCTCCTTCC) and MHC-II DRB exon 2 (F: CGAGTGCCATTTCACCAACG, R: GCTGCACCGTGAAGCTCT) based on sequences of closely related carnivore species. All samples were amplified with a technical replicate and each plate contained a negative control. After initial amplification, samples were purified and normalized using the NGS Normalization 96-well kit (Norgen Biotek Corp., Canada). Libraries were then prepared using the llumina TruSeq DNA Nano Low Throughput Library Prep Kit (Illumina Inc, United States) and the TruSeq DNA Single Indexes Set A and B (Illumina Inc, United States). After checking library quality using the Agilent Bioanalyzer 2100 (Agilent Technologies, United States) together with the High Sensitivity DNA Kit (Agilent Technologies, United States), we removed primer dimers by running the libraries for 3h at 65V on a 1.5% agarose gel, cutting out bands corresponding to the correct amplicon sizes, and finally using either the innuPREP Gel Extraction Kit (Analytik Jena, Germany) or the QIAquick Gel Extraction Kit (Qiagen) according to the manufacturer’s protocol. Libraries were then pooled and sequenced on an Illumina MiSeq® (Illumina Inc, United States) with a MiSeq Reagent Kit v2 (500 cycles) according to the manufacturer’s protocol at the Max Planck Institute for Evolutionary Biology, Plön, Germany. We estimated reproducibility of the genotypes based on the formula [(Number of shared alleles*2)/sum of alleles in replicates] to investigate consistency between the replicates. Reproducibility was 89.4% for the MHC-I exon 2, 96.6% for the MHC-I exon 3, and 97.7% for the MHC-II DRB exon 2. In total, genotypes were established successfully for 321 individuals for the MHC-I exon 2, 285 individuals for the MHC-I exon 3, and 385 individuals for the MHC-II DRB exon 2. Mismatched alleles were kept as putative alleles if they fulfilled the criteria for allele validation described in Schubert et al. (2024). Assigning alleles to loci was not possible, which is why heterozygosity of the MHC could not be estimated. Detailed methods for the characterization of the MHC in banded mongooses are described in Schubert et al. (2024).

#### 2.3.4 Measures of MHC diversity

We estimated MHC diversity for each exon separately using three measures (1) the mean amino acid p-distance, (2) the number of functional alleles and (3) supertype number. Mean amino acid p-distance is defined as the proportion of amino acid sites at which the compared sequences differ. It is calculated by dividing the number of amino acid differences by the total number of amino acid sites compared and represents the functional distance of a genotype. We calculated the mean amino acid p-distance in R version 4.4.0 using the function p.dist of the package phangorn, version 2.12.1 (Schliep 2011). Since we amplified multiple loci and were not able to assign alleles to loci, we cannot include MHC heterozygosity as a measure of MHC diversity. Instead, we used the total number of functional alleles per individual as well as number of supertypes as measures of functional diversity. We determined the number of functional alleles by translating the filtered nucleotide sequences to identify alleles with identical amino acid sequences or stop codons. These analyses were conducted in MEGA X Version 10.0.5 (Kumar et al. 2018). Functional alleles were then assigned to functionally similar supertypes based on hierarchical clustering of the amino acid characterization matrix (Sandberg et al. 1998) estimated based on the positively selected sites (PSS) and k-means as conducted by Winternitz et al. (2015). PSS were identified using the rates of non-synonymous (dN) and synonymous (dS) mutations for each site based on two complimentary methods: FUBAR and MEME. FUBAR (Fast, Unconstrained Bayesian AppRoximation) estimates selection rates based on a Bayesian approach (Murrell et al. 2013). It detects signatures of pervasive selection by assuming constant pressure. As individual sites can experience different levels of positive and negative selection (episodic selection), methods that only detect pervasive selection will miss episodic effects. In contrast, MEME (Mixed Effects Model of Evolution) uses a mixed-effects maximum likelihood approach to determine individual sites with signs of pervasive and episodic selection (Murrell et al. 2012). Hence PSS were either detected by FUBAR (pervasive selection) or MEME (pervasive and episodic selection). For analysis with FUBAR, the posterior probability was set to > 0.9 as our significance threshold, since values above this threshold indicate natural selection. For analyses with MEME, the significance threshold was set to 0.05. Both FUBAR and MEME selection inference was carried out on the Datamonkey server (Weaver et al. 2018, https://www.datamonkey.org/, last accessed: January 18, 2024). Supertypes were determined by assigning sequences to functionally similar supertypes based on k-means and hierarchical clustering of the amino acid characterization matrix (Sandberg et al. 1998) of PSS following the approach of Winternitz et al. (2015).

### 2.4 Statistical analyses

#### 2.4.1 Assessing collinearity among variables

Substantial collinearity among variables can cause problems in model interpretation (including convergence issues, changes in the direction of effects, and inflated standard errors of coefficient estimates) because the effects of collinear predictors cannot be estimated independently (reviewed in Harrison et al. 2018). Thus, we performed correlation analyses to investigate whether MHC diversity measured as mean amino acid p-distance, number of functional alleles per individual, or supertype number, are strongly correlated (correlation coefficient r ≥ 0.7) with each other as well as with neutral genetic diversity (sMLH). To estimate the former, we calculated Pearson’s correlation coefficients in R version 4.4.0 (R Core Team 2023) for correlations between MHC diversity measures. All correlations between MHC measures were highly significant, but the strength of the correlation differed for the exons. Between the MHC diversity measures, there was a strong correlation (r > 0.7; Tab. S4) between MHC-I exon 3 mean amino acid p-distance and number of functional alleles (r = 0.796, p < 0.001), which can be explained by the low allelic diversity of this exon. The correlation between MHC-I exon 3 mean amino acid p-distance and supertype number (r= 0.620, p < 0.001) as well as allele and supertype number (r = 0.695, p < 0.001) for MHC-I exon 3 were comparably high, due to the same cause. However, correlation coefficients for the other MHC exons were often substantially above 0.2 (range -0.268 to 0.573) so we therefore decided to run separate models for each diversity measure.

To test for a relationship between MHC diversity and neutral genetic diversity (sMLH), we ran linear mixed models in R with the package lme4 version 1.1-35.3 (Bates et al. 2017), fitting each MHC diversity measure separately as the dependent variable and sMLH as the independent variable. We also included social group as a random slope in order to account for potential differences in sMLH between the social groups the individuals were born into. In effect, banded mongoose usually stay within their natal pack and with and rarely disperse, resulting in pack relatedness values increasing and thus individual sMLH decreasing over time (Cant et al. 2013). Hence the slope might vary depending on the age and structure of the pack an individual is born in. None of the three MHC diversity measures were strongly correlated with sMLH (r < 0.1; Tab. S1-3, Fig. S1-2), so we included MHC diversity measures together with sMLH as predictor variables in the same models.

#### 2.4.2 Assessing the impact of MHC on fitness

We constructed separate models for each of the three fitness measures by including all potentially biologically relevant independent variables and adjusting model structure if necessary, based on model diagnostics. For each fitness measure, we fitted nine generalized linear mixed models (GLMMs) with the respective fitness measure as the dependent variable. An MHC diversity measure (mean amino acid p-distance, number of functional alleles per individual and supertype number), for each of the three MHC classes and exons (MHC-I exon 2 and 3, and MHC-II) was included as the independent variable and an interaction with sex was fitted. We also included sMLH as an independent variable to account for genome-wide diversity, together with other predictors as described separately in the following sections for the different fitness measures. We removed individuals (n = 2 for MHC-I exon 2, n = 1 for MHC-I exon 3, and n = 3 for MHC-II DRB exon 2) with unrealistic lifespans, such as negative lifespans, as we assumed errors in the life history data. We also removed one individual with an unrealistic sMLH value (sMLH = 0). All analyses were run in R version 4.4.0. The model structure for each of the 27 models can be found in Table S5 in the supplementary material.

##### 2.4.2.1 Pup survival

We ran models analyzing the association between MHC diversity and pup survival using the glmer function of the lme4 package version 1.1-35.3 (Bates et al. 2017). As the dependent variable, we used survival to nutritional independence (90 days of age) as a binomial response (with 1 = survived, and 0 = died). As independent variables, we used one of the measures of MHC diversity in interaction with sex and sMLH as a previous study on banded mongooses found higher pup survival for offspring originating from extra-group matings, which are more heterozygous (Nichols et al. 2015). We also included the amount of rainfall within 30 days prior births (log-transformed) as an independent variable, as this has been observed to be linked to early life survival in banded mongooses (Nichols et al. 2015; Sanderson et al. 2015). For individual MHC allele number, we included a quadratic term to test for nonlinear effects on pup survival; this was ultimately removed as it did not significantly improve the explained variation. Model fit was tested by simulating the residuals and checking for zero inflation and overdispersion using DHARMa version 0.4.6. (Hartig et al. 2024).

##### 2.4.2.2 Adult survival

We ran Cox proportional hazard models to analyze the potential link between MHC diversity and adult survival using the coxph function of the survival package version 3.6-4 (Terry 2024). In our models, data was right censored for individuals that either survived to the end of the study or emigrated. The total lifespan in days was included as the dependent variable and one of the measures of MHC diversity in interaction with sex and sMLH were included as independent variables. For individual MHC allele number, we included a quadratic term to test for nonlinear effects on adult survival but finally removed it from the model as it did not significantly improve the fit. We tested the proportional hazard assumption by investigating and plotting the Schoenfeld residuals against time using the ggcoxzph command and the ggcoxdiagnostics command with the residual “Schoenfeld” from the survMisc package version 0.5.6 (Dardis 2022) and looked for influential observations using the residuals “dfbeta” and for outliers using “deviance” with the ggcoxdiagnostics command from the package survMisc version 0.5.6.

##### 2.4.2.3 Lifetime reproductive success

We ran the glmmTMB function (family = nbinom1, ziformula = ∼1) from the package glmmTMB version 1.1.9 (Brooks et al. 2017) with the total number of offspring assigned in the pedigree to each individual as the dependent variable. This approach of applying a negative binomial distribution with zero inflation for count data has been frequently used in both phytopathology and medicine (e.g. Prager et al. 2014; Almeida et al. 2016). One of the MHC diversity measures in interaction with sex was included as an independent variable. Furthermore, mean monthly rainfall per lifetime and lifespan itself were also included as independent variables. For individual MHC allele number, we included a quadratic term to test for nonlinear effects on total number of pups produced.

#### 2.4.3 Standardization of effect sizes

Effect sizes were exported from the respective models and standardized using the standardize_parameters function from the effectsize package under version 0.8.9. (Ben-Shachar et al. 2020) using the “refit” function.

#### 2.4.4 Correcting for multiple hypotheses testing

We corrected for multiple testing for each dependent variable investigated (pup survival, adult survival, LRS) using the false discovery rate (FDR) (Benjamini and Hochberg 1995). We used the p.adjust function from the stats package (R Core Team 2023) with “fdr” as the method of choice correcting for nine hypotheses tested for each fitness measurement (three MHC genes tested with three MHC diversity measures each). We solely corrected the relevant or significant effects to investigate whether significance holds to reduce Type II error.

## Results

### 3.1 Genotyping results

Genotyping was successful for 321 individuals for the MHC-I exon 2, 282 individuals for the MHC-I exon 3, and 385 individuals for the MHC-II DRB exon 2. After removing a small number of outliers, the number of sequences per individual ranged from 2 to 11 (median = 6, n = 300) for the MHC-I exon 2, between 1 and 5 (median = 1, n = 264) for the MHC-I exon 3, and between 1 and 6 (median = 2, N = 361) for the MHC-II DRB exon 2. Functional differences, quantified as mean amino acid p-distances, ranged from 0.1447 to 0.333 (median = 0.2566) for the MHC-I exon 2, from 0 (only one allele per individual) to 0.1818 (median = 0) for the MHC-I exon 3, and from 0 (only one allele per individual) to 0.3485 (median = 0.2727) for the MHC-II DRB exon 2. Supertype numbers ranged from 1 to 3 (median = 2) for the MHC-I exon 2, from 1 to 2 (median = 1) for the MHC-I exon 3, and from 1 to 2 (median = 2) for the MHC-II DRB exon 2. Detailed results of MHC characterization analyses can be found in Schubert et al. (2024).

### 3.2 Associations between MHC and sMLH

We found no evidence that any of our MHC diversity measures were significantly positively associated with genetic diversity (sMLH) at microsatellite loci (Tables S1-S3).

### 3.3 Pup survival

Neither MHC-I exon 2 nor MHC-I exon 3 diversity, measured as any of the three MHC diversity measures, significantly affected pup survival (Fig.1, Tab. S6). Similarly, MHC-II DRB exon 2 mean amino acid p-diversity and supertype number did not significantly impact pup survival. However, MHC-II DRB exon 2 individual allele number in interaction with sex showed a significant positive association with pup survival (effect size = 0.470, CI = 0.075 – 0.865, p = 0.020, n = 344), indicating a positive relationship between survival and MHC-II allele number in males and a negative relationship in females (Fig. 2). After FDR correction for multiple hypotheses testing, this relationship became marginally non-significant (p_corrected_ = 0.072).

**Figure 1.**
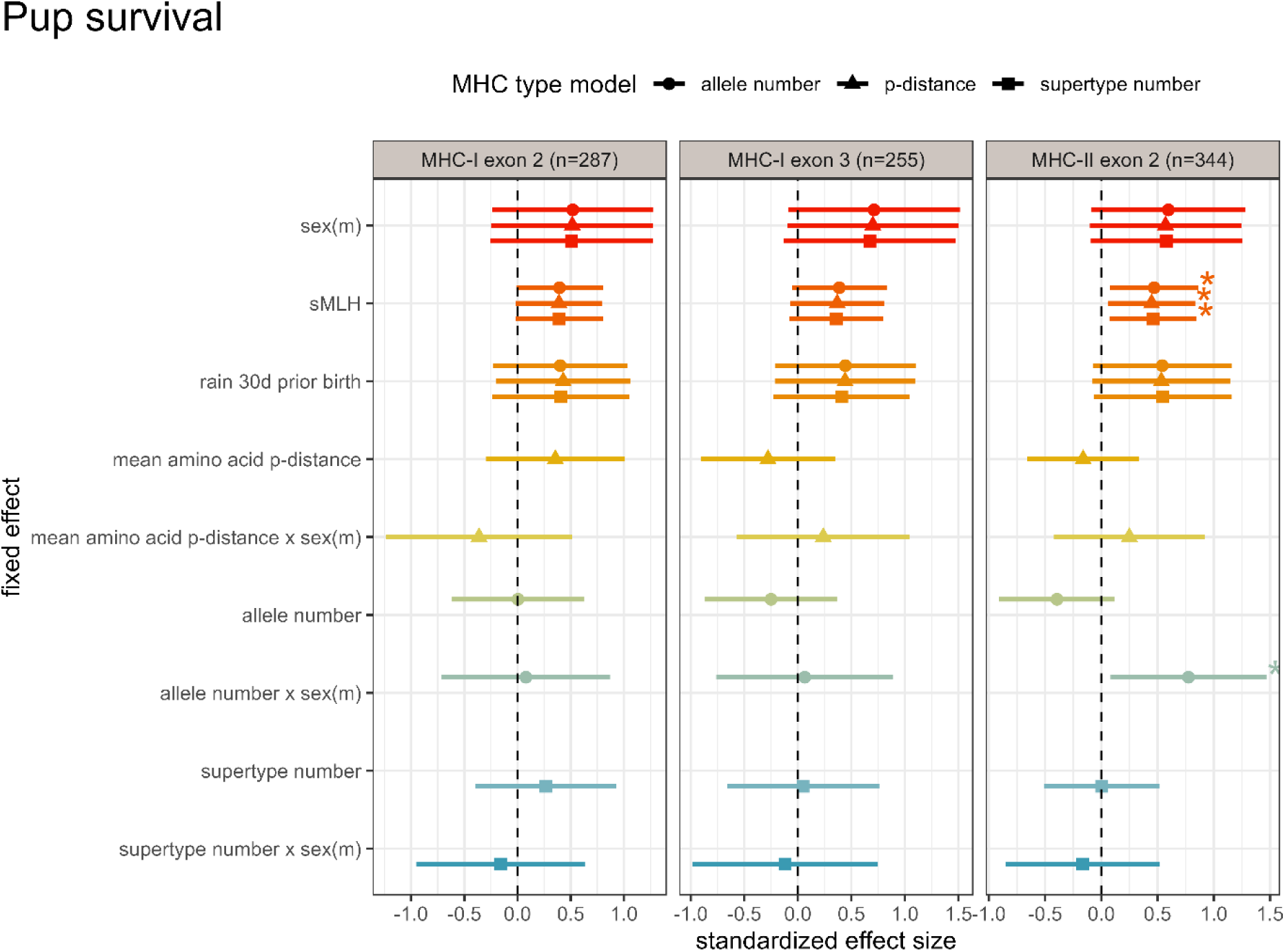
Standardized effect sizes for models of pup survival. Shown are the standardized effect sizes of the models investigating relationships between MHC diversity and pup survival. The effect sizes are shown separately for three different diversity measures (allele number, amino acid p-distance and supertype number) for the three different genes (MHC-I exon 2, 3 and MHC-II DRB exon 2). Asterisks indicate significant variables before FDR correction for multiple testing.

**Figure 2.**
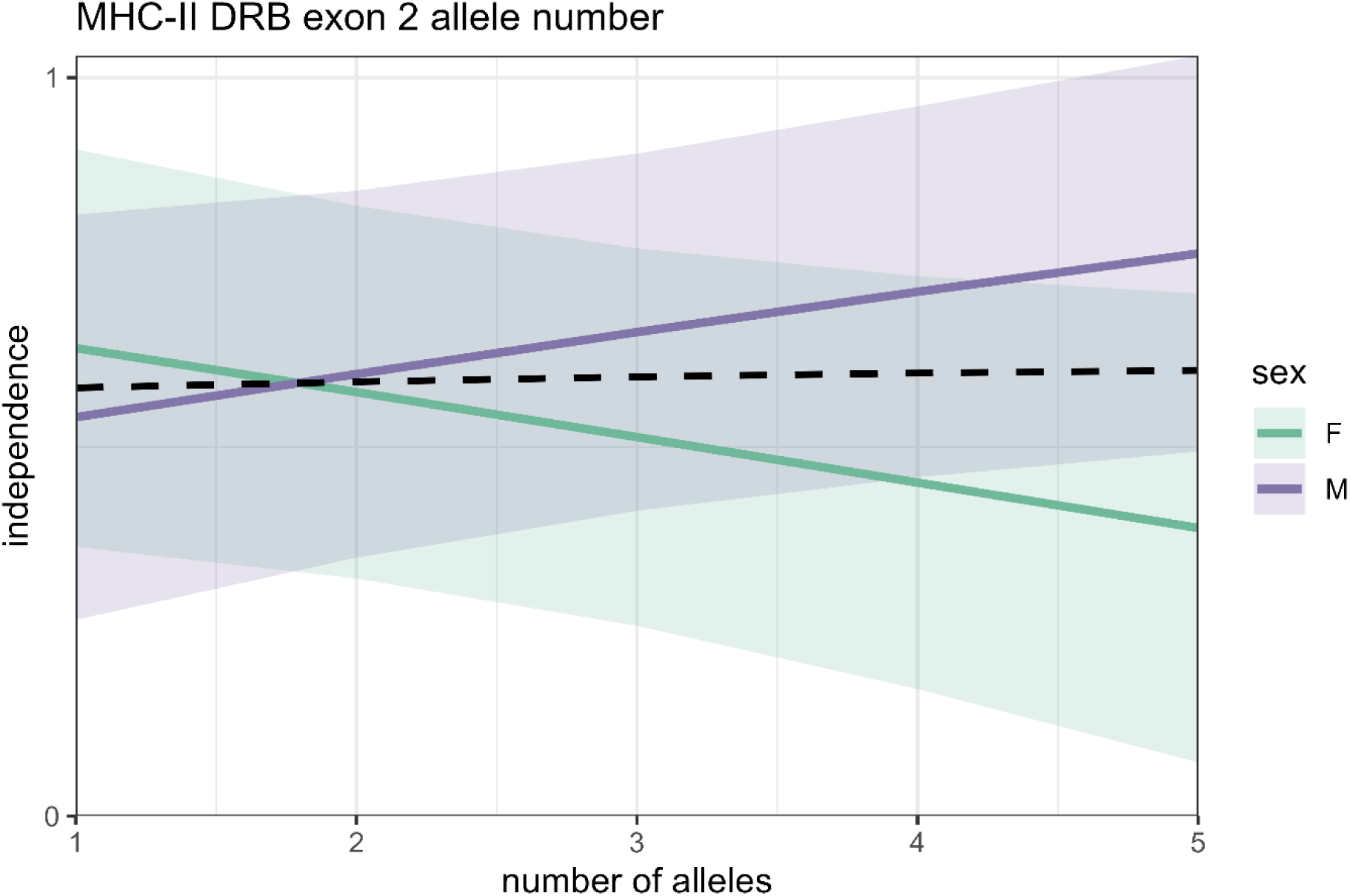
Relationship between individual MHC-II allele number and pup survival. Relationship between individual MHC-II DRB exon 2 allele number and survival to 90 days shown separately for the sexes. Independence is plotted as a binomial variable with 0 indicating that pups did not survive until independence and 1 indicating pup survival until independence. Regression lines are shown for females in green and males in purple, and the dashed black line represents the effect for both sexes combined. Shaded areas represent the corresponding 95% confidence intervals.

In three of our models, neutral genetic diversity (sMLH) had a significant positive effect on pup survival, indicating that inbred pups are less likely to survive to nutritional independence than outbred pups. However, after correcting for multiple hypothesis testing, sMLH lost significance in all of these models (Tab. S6).

### 3.4 Adult survival

None of our MHC diversity measures (nor any other variable that we included in the models) was significantly associated with adult survival (Fig. 3, Tab. S7).

**Figure 3.**
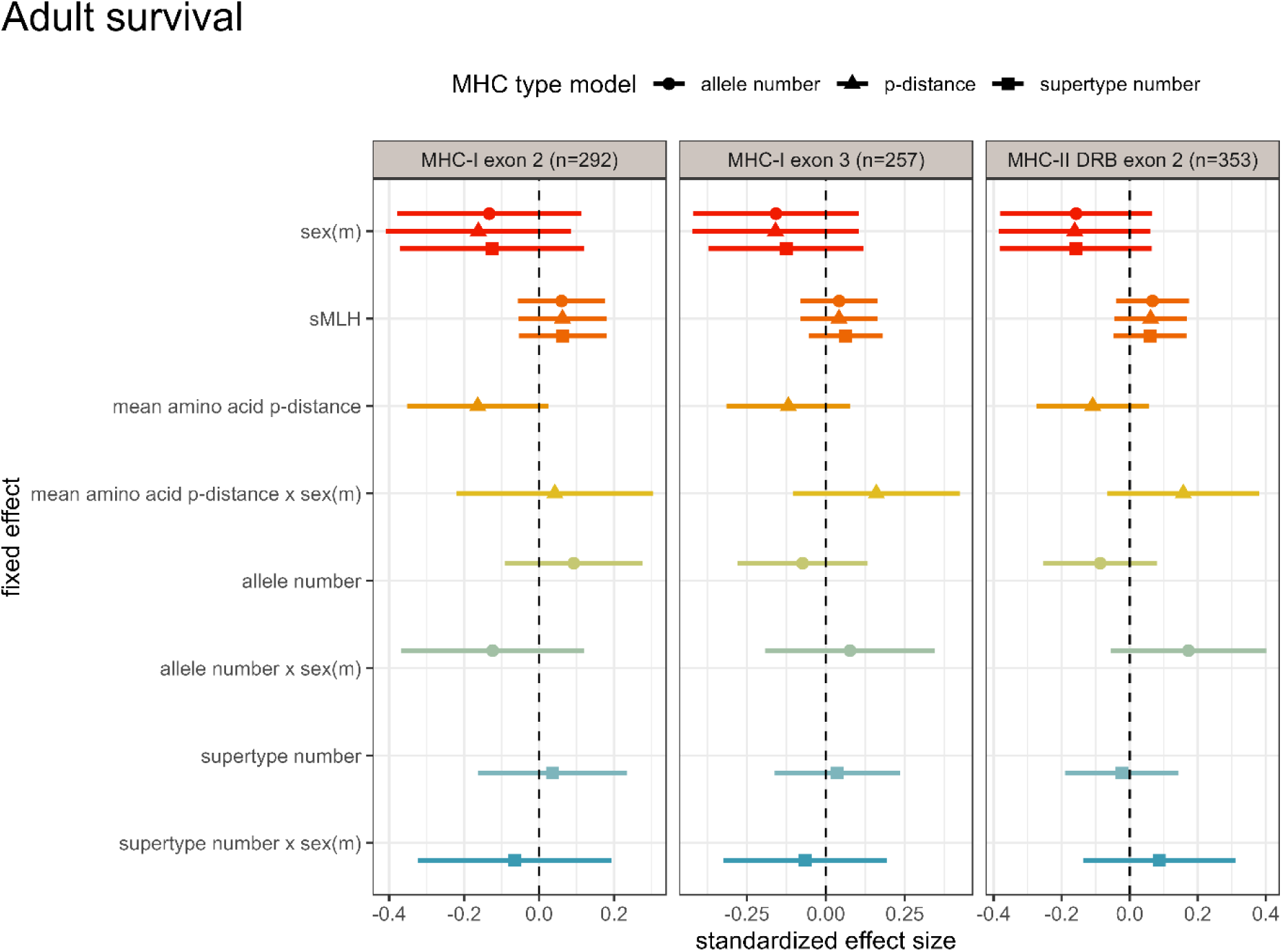
Standardized effect sizes for models of adult survival. Shown are the standardized effect sizes of the models investigating relationships between MHC diversity and adult survival. The effect sizes are shown separately for three different diversity measures (allele number, amino acid p-distance and supertype number) for the three different genes (MHC-I exon 2, 3 and MHC-II DRB exon 2).

### 3.5 Lifetime reproductive success

For four of our models, the interaction between MHC diversity and sex predicted lifetime reproductive success, all of which remained significant after correcting for multiple testing (Fig. 4, Tab. S8). Since sex or its interactions remained significant in five models, we investigated the effects of individual MHC diversity on lifetime reproductive success separately for each sex and observed contrasting directions (Fig. 5+6, Tab. S9).

**Figure 4.**
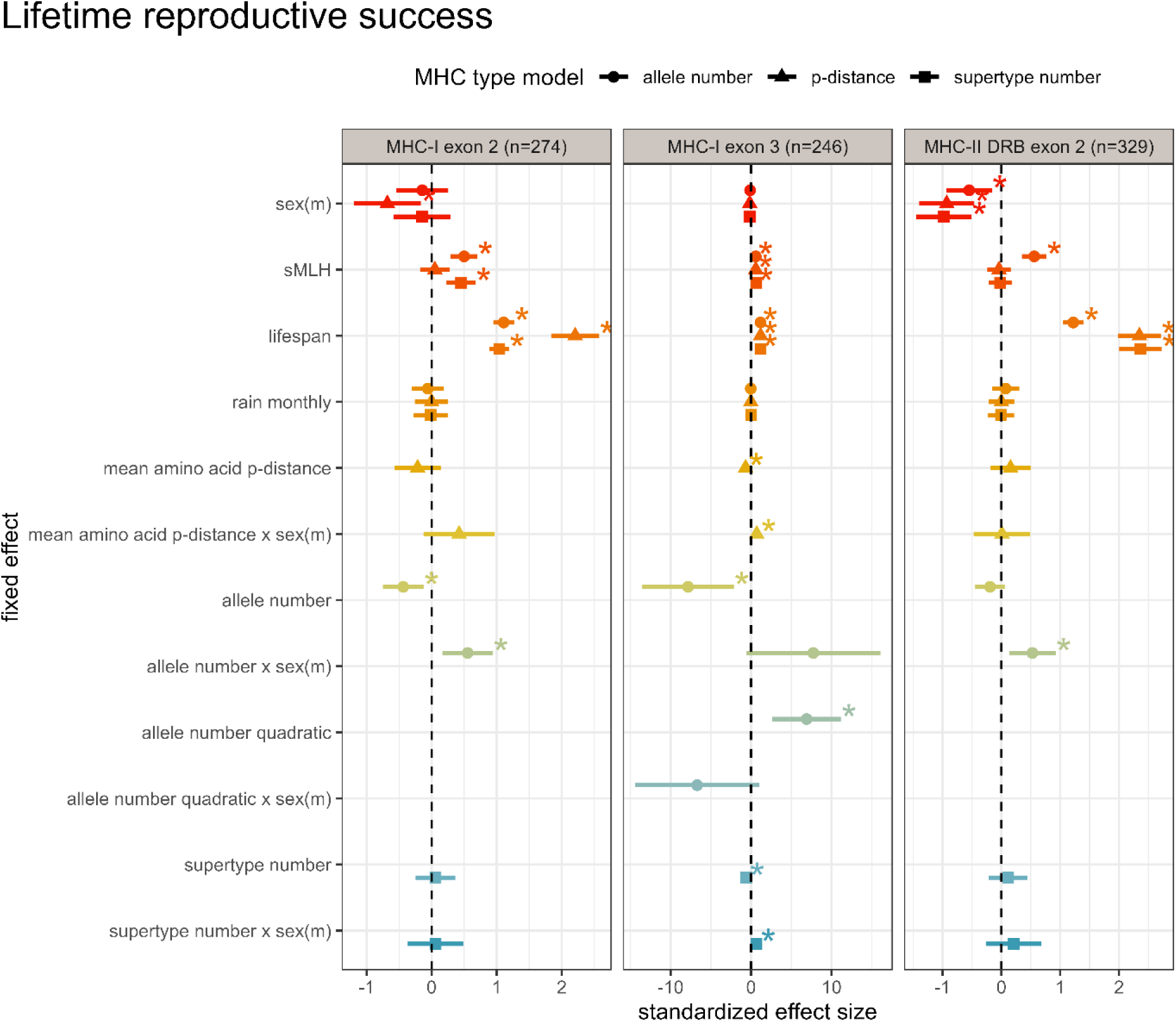
Standardized effect sizes for models investigating lifetime reproductive success. Displayed are the standardized effect sizes of the models investigating links of MHC diversity with lifetime reproductive success. The effect sizes are shown separately for the different diversity measures (allele number, amino acid p-distance and supertype number) for the three different genes (MHC-I exon 2, 3 and MHC-II DRB exon 2). Asterisks indicate significant initial p-values before correcting for multiple hypothesis testing.

**Figure 5.**
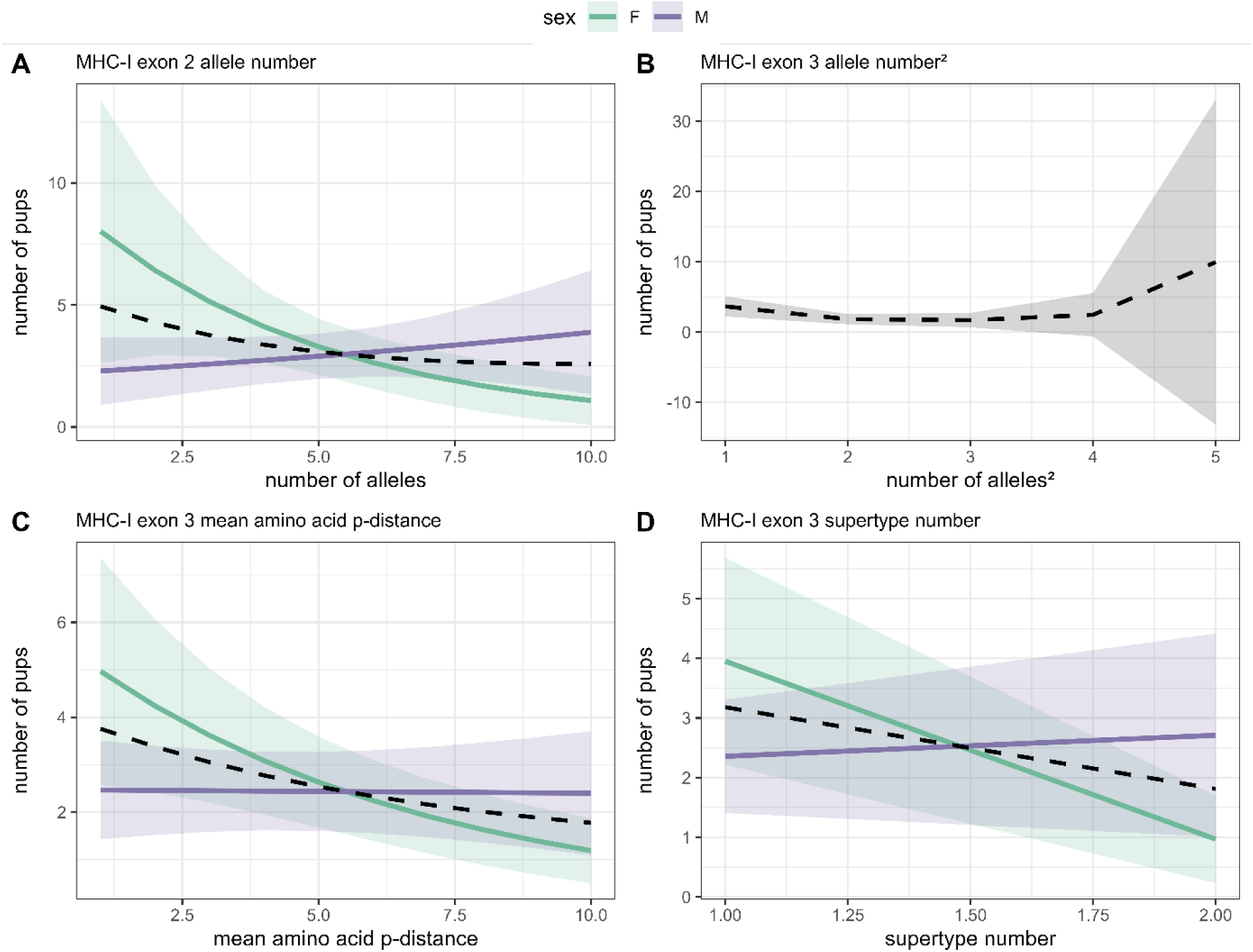
Relationship between individual MHC-I diversity measures and lifetime reproductive success. The investigated variables are individual MHC-I exon 2 allele number separately for the sexes (A), the quadratic number of MHC-I exon 3 alleles (B), individual MHC-I exon 3 mean amino acid p-distance separately for the sexes (C), MHC-I exon 3 supertype number separately for the sexes (D). Lines depict regression lines separately for the sexes (males: purple, females: green) and the dashed black line represents the effect for both sexes combined. Shaded bands show 95% confidence intervals.

For the MHC-I exon 2, female reproductive success declined while male reproductive success increased with increasing allele number (Fig. 5A). The comparison indicated that the slope was significant for females, but not males (Tab. S9). Mean MHC-I exon 2 allele numbers did not differ significantly between the sexes (t-test: mean_female_ = 4.764, mean_male_ = 5.073, CI =-0.779 – 0.162, p = 0.198). Similarly, MHC-II allele number in interaction with sex had a significant effect on lifetime reproductive success in the same fashion (Tab. S9, Fig. 6). However, in contrast to MHC-I exon 2 allele number, the slope was significant for males but not for females (Tab. S9). Similar to the MHC-I exon 2, MHC-II allele number did not differ significantly between the sexes (t-test: mean_female_ = 2.415, mean_male_ = 2.540, CI =-0.321 – 0.072, p = 0.213). The model including MHC-I exon 3 supertype number showed a positive relationship between supertype number and lifetime reproductive success for males (Fig. 5D). In contrast, lifetime reproductive success was negatively linked to individual MHC-I exon 3 allele number for females. Again, the slope for females was significant whereas it was non-significant for males (Tab. S9). Mean MHC-I exon 3 allele numbers did not differ significantly between the sexes (t-test: mean_female_ = 1.185, mean_male_ = 1.276, CI =-0.197 – 0.015, p = 0.093). For MHC-I exon 3 mean amino acid p-distance, the slopes were significantly different for the sexes but both were negative (Fig. 5C). However, only the slope for females was significant. Mean MHC-I exon 3 mean amino acid p-distance did not differ significantly between the sexes (t-test: mean_female_ = 4.084, mean_male_ = 4.622, CI =-1.590 – 0.514, p = 0.315). Moreover, MHC-I exon 3 allele number fitted as a quadratic term had a significant effect on lifetime reproductive success with no effect of sex (Fig. 5B).

**Figure 6.**
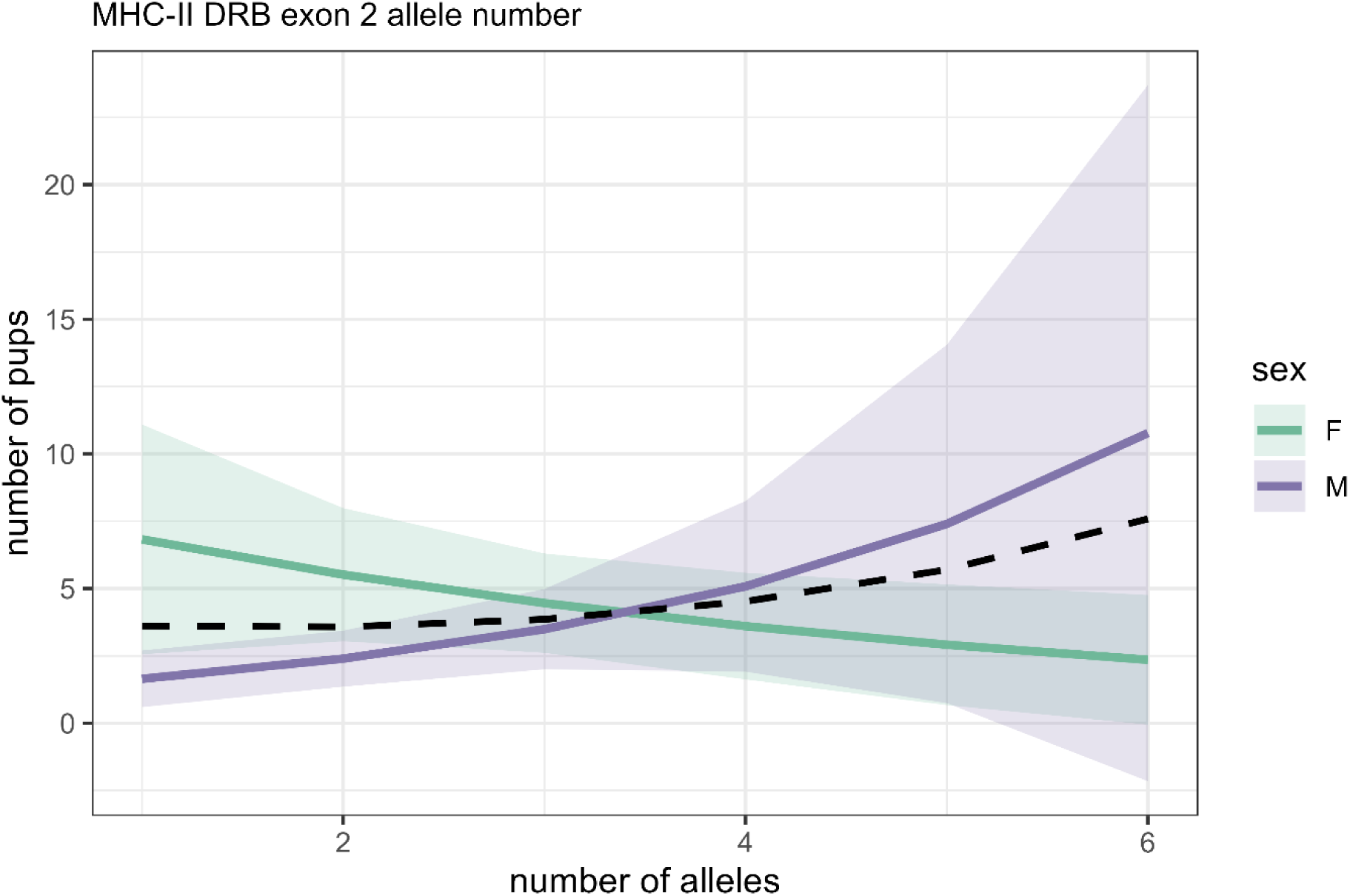
Relationship between individual MHC-II allele number and lifetime reproductive success. **I**ndividual MHC-II DRB exon 2 allele number is shown separately for the sexes. Lines depict regression lines separately for the sexes (males: purple, females: green) and the dashed black line represents the effect for both sexes combined. Shaded bands show 95% confidence intervals.

As expected, lifetime reproductive success was significantly positively associated with lifespan in all models, indicating that individuals who live longer have more offspring. Neutral genetic diversity (sMLH) was also significantly positively correlated with reproductive success in the majority (5 of 8) of our models, indicating that more heterozygous (less inbred) individuals have higher lifetime reproductive success than those with lower heterozygosity. Model estimates can be found in Table S8 and Figure 4.

## Discussion

Investigating effects of MHC diversity on fitness in banded mongooses revealed contrasting sex-specific patterns. Males appeared to benefit from having higher MHC diversity for all three MHC diversity measures: male lifetime reproductive success increased with MHC-I exon 2 allele number, with MHC-I exon 3 supertype number, and with MHC-II DRB exon 2 allele number. By contrast, females showed the opposing pattern, with their reproductive success tending to decline with increased MHC diversity. Hence, our findings lend support to the hypothesis of sexual conflict for MHC diversity.

Our results support previous findings in great reed warblers, which revealed similar relationships between MHC-I diversity and reproductive success, whereby males benefitted from increased MHC diversity, but females showed the opposite pattern (Roved et al. 2018). This pattern likely arises because the sexes differ in their immune responses and in their levels of sex hormones, which in turn have immunosuppressive (testosterone) (Folstad and Karter 1992; Foo et al. 2017) or immune-enhancing (estrogen, progesterone) functions (Foo et al. 2017). Males are therefore predicted to benefit from high MHC diversity through boosting immune responses against pathogens (Folstad and Karter 1992). Females however, may benefit from lower MHC diversity as they are more prone to autoimmune reactions if MHC diversity is high (reviewed in Roved et al. 2017). This difference may in turn cause sexually antagonistic selection.

In banded mongooses, we found no sexual dimorphism in MHC diversity between the sexes, indicating an unresolved sexual conflict. In contrast to other genes, sexual conflict may be difficult to resolve for the MHC, as genes of the MHC are autosomal, being located on chromosomes carried by both sexes (Beck et al. 1999; Kumánovics 2006). This means that MHC alleles from both males and females are combined within an offspring, thus a resolution of the sexual conflict is unlikely (Roved et al. 2018). Nevertheless, it is possible that differences in gene expression could potentially resolve the conflict (Roved et al. 2018). However, as sex differences in MHC gene expression should act to weaken sex-specific relationships between MHC diversity and fitness, our results suggest that it is unlikely that sex-specific differential expression of MHC alleles occurs in banded mongooses. In humans, there is evidence for sex-specific and disease related expression patterns (Mathé et al. 2024). Hence it would be possible that expression differs for different life stages or during parasite infection and that it is specific to the tissue as well. This would be an interesting avenue for future research.

In banded mongooses, several factors relating to the mating system may underly sex-divergent effects of the MHC on fitness. First, during estrus, male group members compete over access to females, with each female being ‘guarded’ by a single male, who attempts to exclude all other males from accessing the female (Cant 2000). The most dominant males usually guard the older, most fecund females (Nichols et al. 2010). Males with low MHC diversity may be more prone to illness, have worse body condition, and therefore be less able to successfully guard females, resulting in lower reproductive success. While this possibility has not yet been tested in banded mongooses, smaller males have been shown to have lower annual reproductive success (Birch et al. 2024) and increased parasite loads compared to larger males (even after controlling for age differences) (Mitchell 2017), which is consistent with this possibility. Furthermore, negative correlations between MHC diversity and parasite loads have been found in other species (e.g. Šimková et al. 2006; Lenz et al. 2009; Meyer-Lucht and Sommer 2009), which would further reinforce the expectation of an effect of MHC-II on fitness through parasite load, although this pattern is not always clear (e.g. Rauch et al. 2006).

Second, female mate choice may play a role in generating sex-dependent effects of MHC diversity as observed in a variety of species (reviewed in Winternitz and Abbate 2022). Moreover, if mate choice in banded mongooses was random with regards to the MHC, we would expect to find a correlation between genome-wide diversity and MHC diversity. We did not observe a significant relationship between any of the three MHC diversity measures and sMLH, which is in line with a study in fur seals that also did not find an association between MHC heterozygosity and sMLH (Tebbe et al. 2024). However, the lack of such a correlation might hint at the possibility of MHC-based mate choice. If female mongooses are like the majority of mammals that prefer males with higher MHC diversity (Kamiya et al. 2014; Winternitz et al. 2017; Winternitz and Abbate 2022), this may contribute to an increase in male reproductive success with increasing MHC diversity through pre-copulatory mate choice. Such a benefit of higher MHC diversity later in life has been observed in macaques, where more MHC-II diverse males have higher reproductive success (Sauermann et al. 2001). While female banded mongooses are guarded by males during estrus, they still appear to exert some choice over mates. Females cannot be forced to mate and are less likely to accept mating attempts by their guard when they are most fertile (Cant 2000). Supporting this, genetic data also reveals that females frequently breed with males other than their guard (Sanderson et al. 2015), which may provide opportunities for MHC-dependent mate choice. Such possibilities should be tested in future studies using careful observation of female behavior. Moreover, apart from female pre-copulatory mate choice, post-copulatory mate-choice mechanisms involving the MHC could result in the observed patterns (Kirby 1970; Amos 1975; Brown 1983) although the immunological and molecular mechanisms behind remain elusive. In banded mongooses, particular effort should be taken to reveal potential post-copulatory mate choice patterns. As litters can be aborted before birth or fail before emergence from the den (Gilchrist 2006a; Gilchrist 2006b), pregnant females should receive ultrasound before birth and close monitoring after birth, to unravel both mating success as well as pregnancy outcomes. Together with pedigree data and behavioral observations, this data could help reveal unsuccessful matings vs. those producing offspring.

If hormonal differences between males and females are the underlying cause for sex-dependent immune defense, MHC-II diversity should be a stronger target for sexually antagonistic selection than MHC-I (Roved et al. 2017). This is because MHC-II affects the humoral (Type 2) response which is the response type that is most strongly impacted by the sex hormones in opposing ways in the sexes (reviewed in Roved et al. 2017). Hence we expected MHC-II diversity to have a greater sex-dependent impact on fitness than MHC-I diversity (Roved et al. 2017). However, while we found that MHC-I diversity had sex-dependent impacts on lifetime reproductive success in four of our nine models (even after correcting for multiple testing), we found a significant effect of MHC-II diversity on lifetime reproductive success for the model investigating MHC-II allele number only. We also found a sex-dependent effect of MHC-II DRB exon 2 allele number on pup survival, but this did not remain significant after correcting for multiple testing. As banded mongooses do not reach sexual maturity until around 1 year of age (Vitikainen et al. 2019), they may show few differences in sex hormones before they reach 90 days of age, explaining why we found little evidence for sex-different responses to MHC diversity on pup survival. Interestingly though, mice and hamsters show a significant difference in serum testosterone levels already 1 to 5 days after birth, so early-life effects may be possible (Pang and Tang 1984). Further studies linking differences in hormonal levels during development to sex-specific immune responses as well as studies linking MHC diversity to disease burdens and fitness may be able to shed light on a potential effect of testosterone-based differences in MHC-diversity and reveal the mechanisms behind it.

MHC-I and MHC-II are both essential for adaptive immunity, but they bind different peptides, which makes each class mainly target specific types of pathogens. MHC-I mostly presents peptides of intracellular origin (Klein 1986), including self-derived peptides as well as peptides originating from microbial pathogens, such as viruses that entered the cell (Rammensee et al. 2013). In contrast, MHC-II predominantly presents peptides that a professional MHC-II carrying and antigen presenting cell has engulfed and these peptides can stem from macroparasites or other pathogens that surround the cell (Neefjes et al. 2011). Hence, the MHC-II should theoretically have a stronger impact on macroparasite load compared to MHC-I. While multiple gastrointestinal and external macroparasites have previously been detected in our study population (Mitchell 2017), the stronger relationship between MHC-I diversity and our fitness measures suggests that microparasites may have stronger impacts on fitness in banded mongooses. In line with this, in Botswana a newly emerged *Mycobacterium tuberculosis* complex pathogen, *M. mungi*, causes significant mortality in banded mongooses and since the outbreak in 1999 93% of the tracked social groups show infections (Alexander et al. 2018), demonstrating a strong impact of microbial pathogens on banded mongoose populations.

In general, our models showed a consistent pattern for the effects of the MHC across the different fitness proxies (Fig. 1, 3 and 4). The direction and magnitude of the standardized effect sizes is similar across models and the contrasting patterns for the sexes was observed across MHC diversity measures. That there were no effects of MHC diversity on adult survival is unlikely to be caused by the model structure. Wells et al. (2018) used a similar model structure and observed sMLH to also have no effect on adult and pup survival, consistent with our findings. Furthermore, we used thorough model diagnostics to ensure the proportional hazard assumption is met and that there are no influential observations or outliers affecting model fit.

The MHC effects varied across the three measures of diversity but showed consistent direction. Differences in the effects of MHC diversity measures, even at the same locus, may result from the three different measures representing slightly different forms of diversity. Mean amino acid p-distance attempts to capture functional diversity of the MHC by using the amino acid sequence to predict the diversity of the PBR and hence the diversity of bound peptides (Wakeland et al. 1990). However, it does not account for the degree of functional change that results from amino acid changes. Because of its three-dimensional structure and binding behavior (reviewed in Lafuente and Reche 2009), it is possible that the amount of amino acid changes does not directly translate into the difference in the binding repertoire. Instead minor mean amino acid p-distances caused by only a small proportion of changes in the amino acid sequence might result in a major increase or decrease in the repertoire of peptides bound by the PBR and significant overlap in the binding repertoire (Rao et al. 2011) Similarly, the number of alleles an individual carries also does not automatically translate into the diversity of the peptides bound. Hence, we additionally used supertype number as a more accurate estimate of the variation in functional MHC diversity of an individual.

When investigating the effect of MHC genotype on fitness, it is important to disentangle the effect of the MHC from effects of genomic background variation. Genes other than the MHC, as well as overall heterozygosity, might impact fitness traits, which could either conceal effects of the MHC or create incorrectly attributed associations (Winternitz 2021; Huang et al. 2021). To mitigate this issue, we included genomic diversity estimated from microsatellite data (sMLH) in our models as a control. We could therefore demonstrate that our MHC diversity measures impacted fitness traits independently of sMLH, rather than due to them being correlated with genomic diversity. Interestingly, our results generally revealed a positive impact of genome-wide diversity on lifetime reproductive success, likely due to inbreeding depression. The findings are in line with a previous study investigating the impact of inbreeding on fitness traits such as yearling body mass and annual reproductive success in male banded mongooses (Wells et al. 2018). In addition, parasite load is negatively associated with heterozygosity in banded mongooses (Mitchell et al. 2017). Our finding of sMLH being a determinant of lifetime reproductive success in the majority of our models suggests that many other genetic loci affect fitness apart from the MHC.

## Conclusion

This study provides evidence for sexually antagonistic selection acting on both MHC classes in a group-living wild mammal with high inbreeding risk resulting in an unresolved conflict. It takes advantage of the exceptional combination of long-term life-history data combined with high throughput sequencing data to reveal contrasting sex-specific effects of MHC diversity on diverse fitness components. By including genetic data for both MHC classes covering the whole PBR region and considering different functional MHC diversity measures, this study is one of the few to use an extensive life-history data set to shed light on the complexity of both classes of the MHC and its impact on fitness traits. Our work highlights the importance of acknowledging and incorporating sex-differences in studies of immunity and fitness measures.

## Acknowledgements

We thank the whole team of the Banded Mongoose Research Project that ensured the continuation of the project and were involved in collecting the invaluable life-history data and genetic samples. We also would like to thank Sven Künzel at the MPI in Plön for Illumina Sequencing.

