## Supplementary material for "Sex-dependent influence of major histocompatibility complex diversity on fitness in a social mammal"

**Table S1 Correlation analysis for mean amino acid p-distance and sMLH.** The table displays the model output for linear mixed models of mean amino acid p-distance explained by sMLH.

| MHC class and exon | Estimate | CI | p-value | n |
|---|---|---|---|---|
| MHC-I exon 2 | 0.01 | -0.01 – 0.02 | 0.450 | 300 |
| MHC-I exon 3 | 0.02 | -0.04 – 0.07 | 0.526 | 264 |
| MHC-II DRB exon 2 | 0.005 | -0.06 – 0.06 | 0.985 | 361 |

**Table S2 Correlation analysis individual MHC allele number and sMLH.** The table displays the model output for linear mixed models of MHC allele number per individual explained by sMLH.

| MHC class and exon | Estimate | CI | p-value | n |
|---|---|---|---|---|
| MHC-I exon 2 | 0.25 | -0.89 – 1.39 | 0.664 | 300 |
| MHC-I exon 3 | 0.32 | -0.12 – 0.77 | 0.153 | 264 |
| MHC-II DRB exon 2 | -0.03 | -0.50 – 0.43 | 0.886 | 361 |

**Table S3 Correlation analysis for supertype number and sMLH.** The table displays the model output for linear mixed models of supertype number explained by sMLH.

| MHC class and exon | Estimate | CI | p-value | n |
|---|---|---|---|---|
| MHC-I exon 2 | 0.05 | -0.19 – 0.28 | 0.694 | 300 |
| MHC-I exon 3 | 0.20 | -0.06 – 0.46 | 0.133 | 264 |
| MHC-II DRB exon 2 | 0.10 | -0.15 – 0.36 | 0.415 | 361 |

**Table S4** Correlations between the MHC diversity measures for the data sets of each model. All of the tested correlations were highly statistically significant ($P \leq 0.001$)

| | Mean amino acid p-distance vs. allele number | Mean amino acid p-distance vs. supertype number | Allele number vs. supertype number |
|---|---|---|---|
| **MHC-I exon 2** | **-0.268*** | **0.238*** | **0.324*** |
| **MHC-I exon 3** | **0.796*** | **0.620*** | **0.695*** |
| **MHC-II DRB exon 2** | **0.573*** | **0.562*** | **0.570*** |

**Table S5 Overview of model structure.** The table shows the model type used to investigate the effect of MHC diversity on the different fitness measures as well as the dependent and independent variables used. An interaction between the MHC diversity measure used in the model (mean amino acid p-diversity, functional allele number or supertype number of one of the genes, MHC-I exon 2 or 3 or MHC-II DRB exon 2) is fitted for each model. Furthermore, the structure of random effects is depicted.

| | *Pup survival* | | | | | | | | | *Adult survival* | | | | | | | | | *Lifetime reproductive success* | | | | | | | | |
|---|---|---|---|---|---|---|---|---|---|---|---|---|---|---|---|---|---|---|---|---|---|---|---|---|---|---|---|
| *Dependent variable* | Survival to nutritional independence (90d of age) | | | | | | | | | Lifespan | | | | | | | | | Total number of offspring | | | | | | | | |
| *Model code* | PS1 | PS2 | PS3 | PS4 | PS5 | PS6 | PS7 | PS8 | PS9 | AS1 | AS2 | AS3 | AS4 | AS5 | AS6 | AS7 | AS8 | AS9 | LRS1 | LRS2 | LRS3 | LRS4 | LRS5 | LRS6 | LRS7 | LRS8 | LRS9 |
| *Model type* | glmer | | | | | | | | | Coxph survival model | | | | | | | | | glmmTMB | | | | | | | | |
| *MHC-I exon 2* | x | | | x | | | x | | | x | | | x | | | x | | | x | | | x | | | x | | |
| *MHC-I exon 3* | | x | | | x | | | x | | | x | | | x | | | x | | | x | | | x | | | x | |
| *MHC-II* | | | x | | | x | | | x | | | x | | | x | | | x | | | x | | | x | | | x |
| *mean amino acid p-distance* | x | x | x | | | | | | | x | x | x | | | | | | | x | x | x | | | | | | |
| *allele number* | | | | x | x | x | | | | | | | x | x | x | | | | | | | x | x | x | | | |
| *(allele number)$^2$* | | | | | | | | | | | | | | | | | | | | | | | x | | | | |
| *suptertype number* | | | | | | | x | x | x | | | | | | | x | x | x | | | | | | | x | x | x |
| *sex* | x | x | x | x | x | x | x | x | x | x | x | x | x | x | x | x | x | x | x | x | x | x | x | x | x | x | x |
| *sMLH* | x | x | x | x | x | x | x | x | x | x | x | x | x | x | x | x | x | x | x | x | x | x | x | x | x | x | x |

| | | | | | | | | | | | | | | | | | | | | | | | | | | | | |
|---|---|---|---|---|---|---|---|---|---|---|---|---|---|---|---|---|---|---|---|---|---|---|---|---|---|---|---|---|
| *Fixed effects* | *MHC diversity variable* sex* | x | x | x | x | x | x | x | x | x | x | x | x | x | x | x | x | x | x | x | x | x | x | x | x | x | x | x |
| | *(allele number)$^2$* sex* | | | | | | | | | | | | | | | | | | | | | | | | x | | | |
| | *rain 30d prior birth* | x | x | x | x | x | x | x | x | x | | | | | | | | | | | | | | | | | | |
| | *rain monthly* | | | | | | | | | | | | | | | | | | | x | x | x | x | x | x | x | x | x |
| | *lifespan* | | | | | | | | | | | | | | | | | | | x | x | x | x | x | x | x | x | x |
| *Random effects* | | (1 \| birth pack/litter) | | | | | | | | | | | | | | | | | | (1\| birth pack) | | | | | | | | |

**Table S6 Model outputs for pup survival.** Summary of results from models JS1 to 9 depicting the relationships of MHC diversity and pup survival. Shown are standardized z-values, upper and lower 95%-confidence intervals, sample size and initial and corrected p-values. P-values were only corrected for relevant/significant effects using FDR correction for multiple testing. Asterisks indicate significant initial p-values (p) and p-values that remained significant after correcting for multiple testing ($p_{corr}$).

| *Model code* | *MHC class and exon* | *Fixed effects* | *Effect size* | *n* | *lower 0.95* | *upper 0.95* | *P* | $p_{corr}$ |
|---|---|---|---|---|---|---|---|---|
| *JS1* | MHC-I exon 2 | mean amino acid p-distance | 0.355 | 287 | -0.296 | 1.007 | 0.281 | |
| | | Sex | 0.515 | | -0.246 | 1.276 | 0.355 | |
| | | rain 30d prior birth | 0.429 | | -0.201 | 1.059 | 0.182 | |
| | | sMLH | 0.390 | | -0.016 | 0.795 | 0.060 | 0.092 |
| | | mean amino acid p-distance*sex | -0.361 | | -1.237 | 0.514 | 0.414 | 0.878 |
| *JS2* | MHC-I exon 3 | mean amino acid p-distance | -0.279 | 255 | -0.907 | 0.349 | 0.384 | |
| | | Sex | 0.700 | | -0.100 | 1.501 | 0.330 | |
| | | rain 30d prior birth | 0.441 | | -0.215 | 1.098 | 0.188 | |
| | | sMLH | 0.368 | | -0.070 | 0.806 | 0.100 | 0.107 |
| | | mean amino acid p-distance*sex | 0.238 | | -0.572 | 1.047 | 0.565 | 0.878 |
| *JS3* | MHC-II DRB exon 2 | mean amino acid p-distance | -0.162 | 344 | -0.659 | 0.335 | 0.524 | |
| | | Sex | 0.571 | | -0.103 | 1.244 | 0.958 | |
| | | rain 30d prior birth | 0.534 | | -0.079 | 1.146 | 0.088 | |
| | | sMLH | 0.447 | | 0.059 | 0.834 | **0.024*** | 0.072 |
| | | mean amino acid p-distance*sex | 0.249 | | -0.422 | 0.920 | 0.466 | 0.878 |
| *JS4* | MHC-I exon 2 | allele number | 0.005 | 287 | -0.619 | 0.628 | 0.989 | |
| | | Sex | 0.519 | | -0.238 | 1.276 | 0.823 | |
| | | rain 30d prior birth | 0.401 | | -0.232 | 1.035 | 0.214 | |
| | | sMLH | 0.395 | | -0.012 | 0.802 | 0.057 | 0.0921 |
| | | allele number*sex | 0.079 | | -0.712 | 0.870 | 0.845 | 0.878 |
| *JS5* | MHC-I exon 3 | allele number | -0.250 | 255 | -0.868 | 0.368 | 0.428 | |
| | | Sex | 0.712 | | -0.091 | 1.514 | 0.547 | |
| | | rain 30d prior birth | 0.444 | | -0.214 | 1.102 | 0.186 | |
| | | sMLH | 0.387 | | -0.056 | 0.831 | 0.087 | 0.107 |
| | | allele number*sex | 0.065 | | -0.760 | 0.890 | 0.878 | 0.878 |
| *JS6* | MHC-II DRB exon 2 | allele number | -0.394 | 344 | -0.909 | 0.120 | 0.133 | |
| | | Sex | 0.596 | | -0.087 | 1.279 | 0.126 | |
| | | rain 30d prior birth | 0.542 | | -0.075 | 1.159 | 0.085 | |
| | | sMLH | 0.470 | | 0.075 | 0.865 | **0.020*** | 0.072 |
| | | allele number*sex | 0.775 | | 0.081 | 1.469 | **0.029*** | 0.258 |
| *JS7* | MHC-I exon 2 | supertype number | 0.793 | 287 | 0.589 | 0.721 | 0.434 | |
| | | Sex | 0.578 | | 0.431 | 0.601 | 0.541 | |
| | | rain 30d prior birth | 2.226 | | 0.971 | 0.982 | 0.215 | |
| | | sMLH | 1.854 | | 0.940 | 0.962 | 0.061 | 0.092 |
| | | supertype number*sex | -0.356 | | -0.440 | -0.235 | 0.695 | 0.878 |
| *JS8* | MHC-I exon 3 | supertype number | 0.024 | 255 | -0.099 | 0.146 | 0.892 | |
| | | Sex | 0.696 | | 0.517 | 0.675 | 0.435 | |
| | | rain 30d prior birth | 1.895 | | 0.944 | 0.965 | 0.208 | |
| | | sMLH | 1.609 | | 0.902 | 0.939 | 0.107 | 0.107 |

| | | | | | | | | |
|---|---|---|---|---|---|---|---|---|
| | | supertype number*sex | -0.157 | | -0.273 | -0.034 | 0.787 | 0.878 |
| JS9 | MHC-II DRB exon 2 | supertype number | 0.037 | 344 | -0.069 | 0.142 | 0.987 | |
| | | Sex | 0.995 | | 0.711 | 0.801 | 0.353 | |
| | | rain 30d prior birth | 2.838 | | 0.992 | 0.994 | 0.081 | |
| | | sMLH | 2.303 | | 0.976 | 0.984 | **0.020*** | 0.072 |
| | | supertype number*sex | -0.531 | | -0.563 | -0.401 | 0.638 | 0.878 |

**Table S7 Model outputs for adult survival.** Summary of results from models AS1 to 9 depicting the relationships of MHC diversity and adult survival. Shown are standardized effect sizes, upper and lower 95%-confidence intervals, sample size.

| *Model code* | *MHC class and exon* | *Fixed effects* | *Effect size* | *lower 0.95* | *upper 0.95* | *n* | *p* |
|---|---|---|---|---|---|---|---|
| *AS1* | MHC-I exon 2 | mean amino acid p-distance | -0.164 | -0.352 | 0.025 | 292 | 0.089 |
| | | sex | -0.162 | -0.409 | 0.085 | | 0.674 |
| | | sMLH | 0.062 | -0.055 | 0.180 | | 0.299 |
| | | mean amino acid p-distance*sex | 0.042 | -0.221 | 0.305 | | 0.754 |
| *AS2* | MHC-I exon 3 | mean amino acid p-distance | -0.119 | -0.314 | 0.077 | 257 | 0.235 |
| | | sex | -0.159 | -0.423 | 0.104 | | 0.094 |
| | | sMLH | 0.042 | -0.080 | 0.163 | | 0.502 |
| | | mean amino acid p-distance*sex | 0.160 | -0.104 | 0.424 | | 0.236 |
| *AS3* | MHC-II DRB exon 2 | mean amino acid p-distance | -0.109 | -0.274 | 0.056 | 353 | 0.197 |
| | | sex | -0.162 | -0.384 | 0.061 | | 0.062 |
| | | sMLH | 0.061 | -0.045 | 0.168 | | 0.255 |
| | | mean amino acid p-distance*sex | 0.158 | -0.066 | 0.381 | | 0.167 |
| *AS4* | MHC-I exon 2 | allele number | 0.093 | -0.092 | 0.277 | 292 | 0.324 |
| | | sex | -0.133 | -0.379 | 0.113 | | 0.541 |
| | | sMLH | 0.060 | -0.057 | 0.177 | | 0.313 |
| | | allele number*sex | -0.124 | -0.368 | 0.120 | | 0.319 |
| *AS5* | MHC-I exon 3 | allele number | -0.074 | -0.279 | 0.132 | 257 | 0.483 |
| | | sex | -0.158 | -0.421 | 0.105 | | 0.311 |
| | | sMLH | 0.042 | -0.080 | 0.164 | | 0.499 |
| | | allele number*sex | 0.077 | -0.192 | 0.345 | | 0.577 |
| *AS6* | MHC-II DRB exon 2 | allele number | -0.087 | -0.254 | 0.081 | 353 | 0.310 |
| | | sex | -0.158 | -0.381 | 0.066 | | 0.062 |
| | | sMLH | 0.067 | -0.041 | 0.175 | | 0.222 |
| | | allele number*sex | 0.173 | -0.056 | 0.401 | | 0.139 |
| *AS7* | MHC-I exon 2 | supertype number | 0.036 | -0.162 | 0.235 | 292 | 0.722 |
| | | sex | -0.126 | -0.371 | 0.120 | | 0.754 |
| | | sMLH | 0.063 | -0.054 | 0.180 | | 0.291 |
| | | supertype number*sex | -0.065 | -0.324 | 0.194 | | 0.622 |
| *AS8* | MHC-I exon 3 | supertype number | 0.036 | -0.162 | 0.235 | 257 | 0.537 |
| | | sex | -0.126 | -0.371 | 0.120 | | 0.371 |
| | | sMLH | 0.063 | -0.054 | 0.180 | | 0.503 |

| | | | | | | | |
|---|---|---|---|---|---|---|---|
| | | supertype number*sex | -0.065 | -0.324 | 0.194 | | 0.584 |
| *AS9* | MHC-II DRB exon 2 | supertype number | -0.023 | -0.190 | 0.144 | 353 | 0.786 |
| | | sex | -0.158 | -0.381 | 0.064 | | 0.256 |
| | | sMLH | 0.060 | -0.047 | 0.167 | | 0.275 |
| | | supertype number*sex | 0.087 | -0.137 | 0.310 | | 0.448 |

**Table S8 Model output for lifetime reproductive success.** Summary of results from models LRS1 to 9 depicting the relationships of MHC diversity and lifetime reproductive success. Shown are z-values, upper and lower 95%-confidence intervals, sample size initial and corrected and p-values. P-values were only corrected for relevant/significant effects using FDR correction for multiple testing. Asterisks indicate significant variables before correcting for multiple testing (p) and after multiple comparison correction ($p_{corr}$).

| *Model code* | *MHC class and exon* | *Fixed effects* | *z-value* | *N* | *lower 0,95* | *upper 0,95* | *p* | $p_{corr}$ |
|---|---|---|---|---|---|---|---|---|
| *LRS1* | MHC-I exon 2 | mean amino acid p-distance | -0.2166 | 274 | -0.57334 | 0.140149 | 0.2431 | 0.3510 |
| | | sex | -0.6817 | | -1.19618 | -0.16713 | 0.0846 | 0.1269 |
| | | sMLH | 0.0483 | | -0.17865 | 0.275307 | 0.6764 | 0.7797 |
| | | rain monthly | 2.2074 | | 1.837218 | 2.577559 | 0.9862 | |
| | | lifespan | -0.0023 | | -0.25967 | 0.255145 | **$<2^{-16*}$** | **$2^{-16*}$** |
| | | mean amino acid p-distance*sex | 0.4221 | | -0.12625 | 0.97053 | 0.1314 | 0.1971 |
| *LRS2* | MHC-I exon 3 | mean amino acid p-distance | -0.6856 | 246 | -1.03738 | -0.33371 | **0.0001*** | **0.0012*** |
| | | sex | -0.1684 | | -0.57986 | 0.243145 | **0.0119*** | **0.0268*** |
| | | sMLH | 0.5741 | | 0.36126 | 0.786863 | **$1.24^{-07*}$** | **$4.62^{-07*}$** |
| | | rain monthly | 1.1822 | | 0.990106 | 1.374289 | 0.7845 | |
| | | lifespan | -0.0366 | | -0.29916 | 0.225897 | **$<2^{-16*}$** | **$2^{-16*}$** |
| | | mean amino acid p-distance*sex | 0.6976 | | 0.204511 | 1.19059 | **0.0056*** | **0.0203*** |
| *LRS3* | MHC-II DRB exon 2 | mean amino acid p-distance | 0.1565 | 329 | -0.18486 | 0.497899 | 0.3689 | 0.4743 |
| | | sex | -0.9289 | | -1.39684 | -0.46086 | 0.1041 | 0.1338 |
| | | sMLH | -0.0401 | | -0.23927 | 0.15907 | 0.6931 | 0.7797 |
| | | rain monthly | 0.0005 | | -0.21945 | 0.220521 | 0.9962 | |
| | | lifespan | 2.3475 | | 1.98241 | 2.712525 | **$<2^{-16*}$** | **$2^{-16*}$** |
| | | mean amino acid p-distance*sex | 0.0106 | | -0.47153 | 0.492684 | 0.9657 | 0.9657 |
| *LRS4* | MHC-I exon 2 | allele number | -0.4383 | 274 | -0.751 | -0.12575 | **0.0060*** | **0.0159*** |
| | | sex | -0.1458 | | -0.5472 | 0.255684 | **0.0038*** | **0.0170*** |
| | | sMLH | 0.4985 | | 0.294054 | 0.702932 | **$1.76^{-06*}$** | **$3.168e^{-06*}$** |
| | | rain monthly | -0.0587 | | -0.30543 | 0.188103 | 0.6413 | |
| | | lifespan | 1.1077 | | 0.947341 | 1.268087 | **$<2^{-16*}$** | **$2e^{-16*}$** |
| | | allele number*sex | 0.5535 | | 0.164719 | 0.942247 | **0.0053*** | **0.0203*** |
| *LRS5* | MHC-I exon 3 | allele number | -7.8419 | 246 | -13.5481 | -2.13574 | **0.0071*** | **0.0159*** |
| | | allele number quadratic | 6.8959 | | 2.628143 | 11.16366 | **0.0015*** | |
| | | sex | -0.1296 | | -0.54691 | 0.287733 | 0.5428 | 0.6106 |
| | | sMLH | 0.5741 | | 0.355141 | 0.793105 | **$2.77^{-07*}$** | **$6.2325^{-07*}$** |
| | | rain monthly | -0.0449 | | -0.32853 | 0.238804 | 0.7566 | |
| | | lifespan | 1.1669 | | 0.960789 | 1.373052 | **$<2^{-16*}$** | **$2^{-16*}$** |
| | | allele number*sex | 7.7370 | | -0.60583 | 16.07985 | 0.0691 | 0.1244 |
| | | allele number quadratic*sex | -6.7098 | | -14.4206 | 1.00108 | 0.0881 | |
| *LRS6* | | allele number | -0.1911 | 329 | -0.44529 | 0.063005 | 0.1405 | 0.2528 |
| | | sex | -0.5477 | | -0.93481 | -0.16049 | **0.0012*** | **0.0106*** |
| | | sMLH | 0.5580 | | 0.349575 | 0.766396 | **$1.54^{-07*}$** | **$4.62^{-07*}$** |

| | | | | | | | | |
|---|---|---|---|---|---|---|---|---|
| | MHC-II DRB exon 2 | rain monthly | 0.0728 | | -0.1597 | 0.305285 | 0.5395 | |
| | | lifespan | 1.2234 | | 1.050603 | 1.396271 | **<2$^{-1}$*** | **2$^{-16}$*** |
| | | allele number*sex | 0.5285 | | 0.131762 | 0.92519 | **0.0090** | **0.0203*** |
| *LRS7* | MHC-I exon 2 | supertype number | 0.0572 | 274 | -0.24853 | 0.363027 | 0.7137 | 0.7137 |
| | | sex | -0.1509 | | -0.58842 | 0.286702 | 0.7043 | 0.7034 |
| | | sMLH | 0.4515 | | 0.228823 | 0.674184 | **7.07$^{-05}$*** | **1.0605$^{-04}$*** |
| | | rain monthly | -0.0168 | | -0.27929 | 0.245793 | 0.9005 | |
| | | lifespan | 1.0390 | | 0.88596 | 1.192122 | **<2$^{-16}$*** | **2$^{-16}$*** |
| | | supertype number*sex | 0.0582 | | -0.36812 | 0.484573 | 0.7890 | 0.8876 |
| *LRS8* | MHC-I exon 3 | supertype number | -0.5948 | 246 | -0.94337 | -0.24631 | **0.0008*** | **0.0025*** |
| | | sex | -0.1583 | | -0.57931 | 0.262709 | **0.0058*** | **0.0175*** |
| | | sMLH | 0.6399 | | 0.431891 | 0.847875 | **1.64$^{-9}$*** | **1.476$^{-08}$*** |
| | | rain monthly | -0.0044 | | -0.26243 | 0.253731 | 0.9737 | |
| | | lifespan | 1.1741 | | 0.976259 | 1.372025 | **<2$^{-16}$*** | **2$^{-16}$*** |
| | | supertype number*sex | 0.6539 | | 0.1641 | 1.1436 | **0.0089*** | **0.0203*** |
| *LRS9* | MHC-II DRB exon 2 | supertype number | 0.1144 | 329 | -0.2166 | 0.4454 | 0.4981 | 0.7137 |
| | | sex | -0.9769 | | -1.4458 | -0.5080 | **0.0439*** | 0.07902 |
| | | sMLH | -0.0198 | | -0.2171 | 0.1776 | 0.8444 | 0.8444 |
| | | rain monthly | -0.0037 | | -0.2257 | 0.2182 | 0.9737 | |
| | | lifespan | 2.3644 | | 2.0032 | 2.7256 | **<2$^{-16}$*** | **2$^{-16}$*** |
| | | supertype number*sex | 0.2114 | | -0.2592 | 0.6820 | 0.3785 | 0.4866 |

**Table S9 Comparison of MHC diversity on the sexes.** The table shows the slopes of MHC diversity effects separately for the sexes for the different models on lifetime reproductive success. Effects with confidence intervals that do not intersect zero are in bold.

| Model | MHC class & exon | | sex | B | SE | CI | n |
|---|---|---|---|---|---|---|---|
| LRS2 | MHC-I exon 3 | mean amino acid p-distance | **F** | **-0.15869** | **0.0432** | **-0.2434 – -0.0740** | **118** |
| | | | M | -0.00304 | 0.0385 | -0.0785 – 0.0724 | 126 |
| LRS4 | MHC-I exon 2 | allele number | **F** | **-0.2226** | **0.0810** | **-0.3813 – -0.0639** | **123** |
| | | | M | 0.0584 | 0.0618 | -0.0626 – 0.1795 | 150 |
| LRS6 | MHC-II DRB exon 2 | allele number | F | -0.213 | 0.144 | -0.4962 – 0.0702 | 141 |
| | | | **M** | **0.376** | **0.166** | **0.0498 – 0.0720** | **186** |
| LRS8 | MHC-I exon 3 | supertype number | **F** | **-1.41** | **0.421** | **-2.23 – -0.583** | **118** |
| | | | M | 0.140 | 0.362 | -0.570 – 0.849 | 126 |

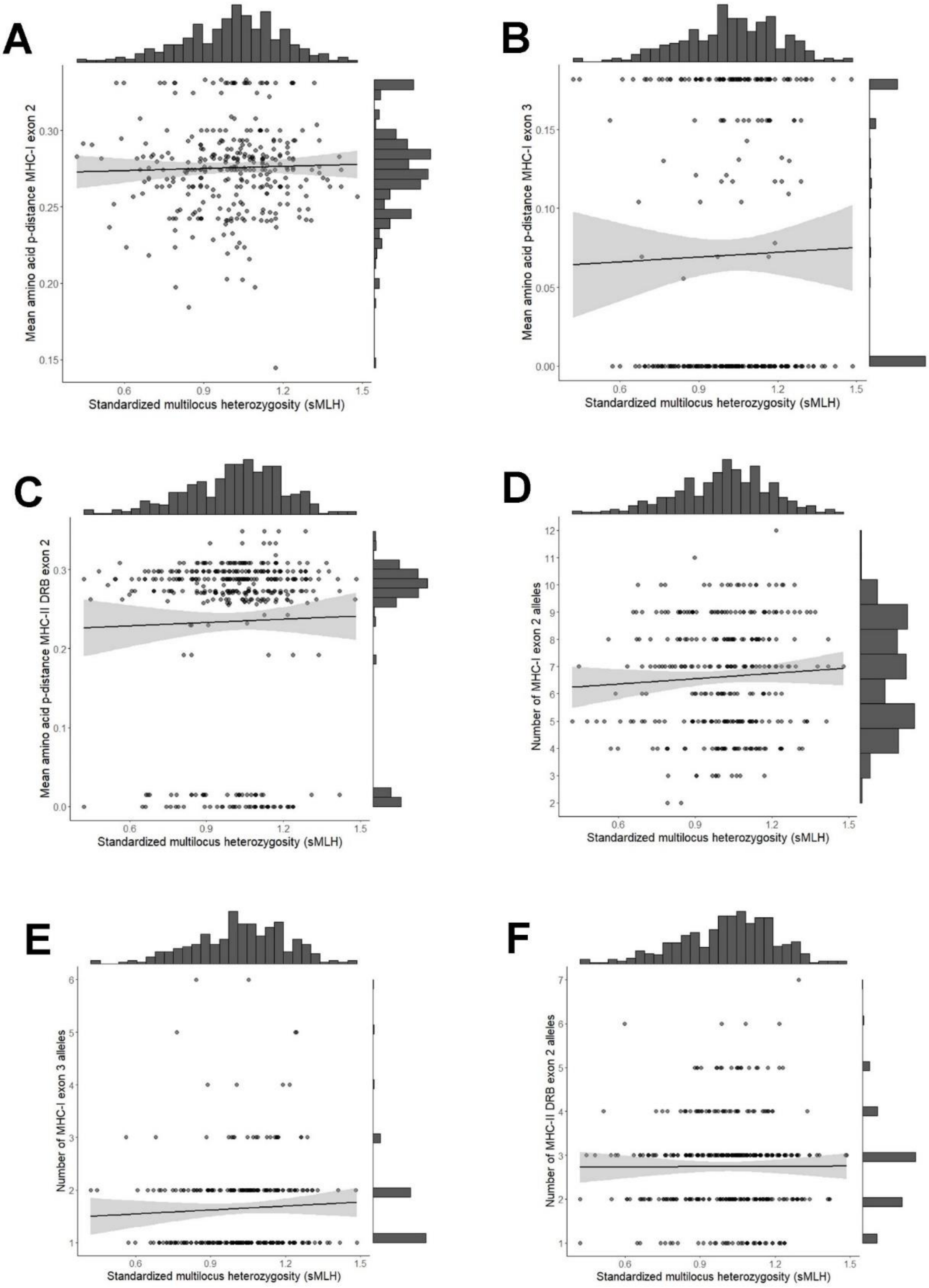

**Figure S1 Correlation between MHC diversity measures and sMLH.** The graphs show the raw data of mean amino acid p-distance (A-C) and individual allele number (D-F) for the three different exons plotted against the sMLH. Marginal histograms visualize the distribution of these values within the sample.

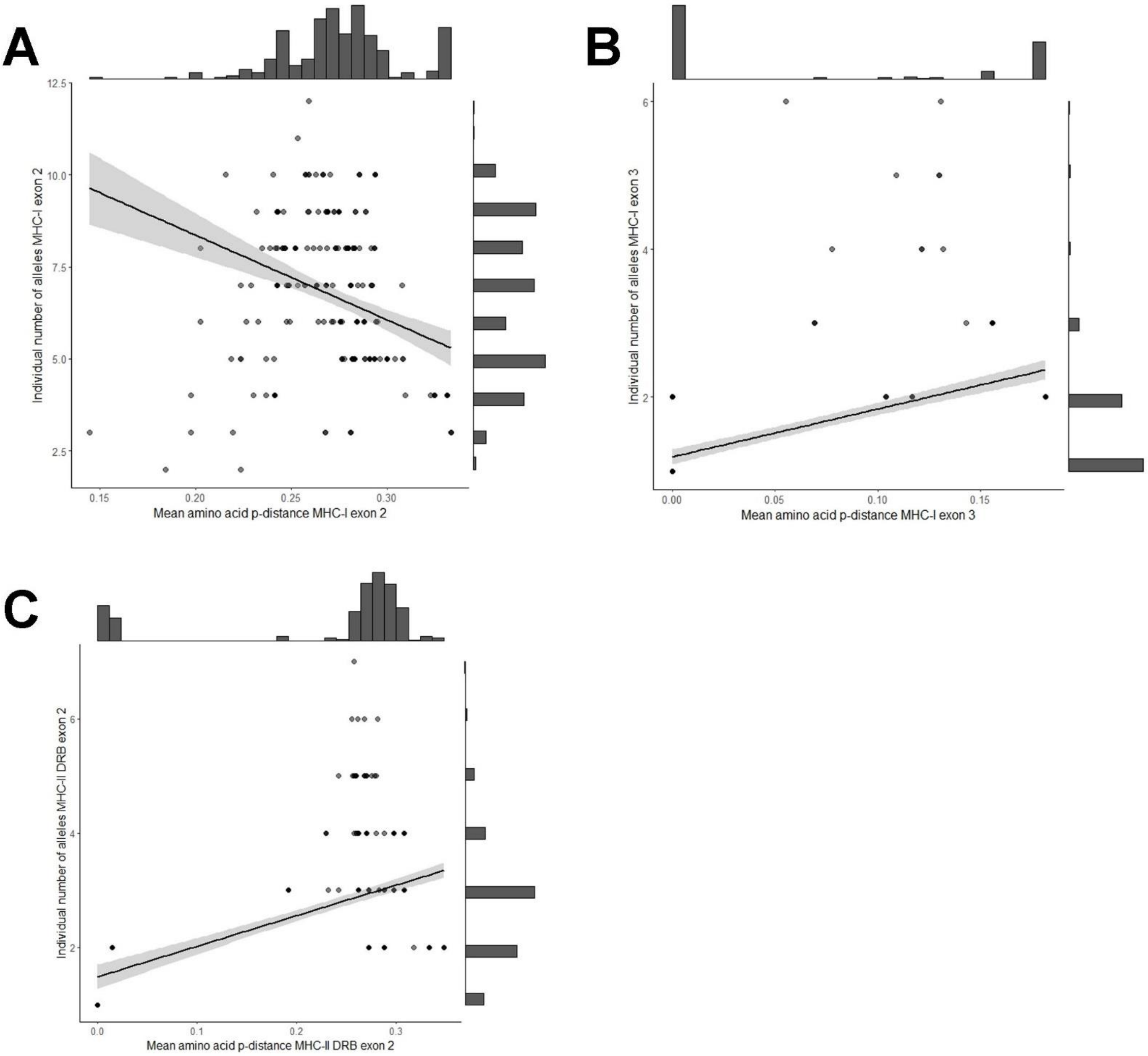

**Figure S2 Correlation between MHC diversity measures.** The graphs show the raw data of mean amino acid p-distance plotted against individual allele number for MHC-I exon 2 (A), MHC-I exon 3 (B), and MHC-II DRB exon 2 (C). Marginal histograms visualize the distribution of these values within the sample.
